# TCRi: Information theoretic metrics for single cell RNA and TCR sequencing in cancer

**DOI:** 10.1101/2022.10.01.510457

**Authors:** Nicholas Ceglia, Zachary M. Sethna, Yuval Elhanati, Bharat Burman, Andrew Chow, Dmitriy Zamarin, Susan DeWolf, Sanam Shahid, Viktoria Bojilova, Nicole Rusk, Vinod P. Balachandran, Andrew McPherson, Sohrab P. Shah, Benjamin D. Greenbaum

## Abstract

Single-cell T cell repertoire sequencing can pair both T cell receptor (TCR) and gene expression sequence data, providing an enriched view of T cell behavior. This powerful tool can identify and characterize specific clonotypes and phenotypes as well as track their changes in response to therapy, such as immune checkpoint blockade (ICB). We present a novel information theoretic framework called TCRi for characterizing single cell T cell repertoires by formalizing the relationship between clonotype and phenotype in a joint probability distribution. Our strategy allows for the identification of subpopulations of T cells and jointly quantifies their TCR and expression profiles in response to stimuli, in addition the framework tracks the phenotypic changes in individual T cell clones over time. We applied this framework to four datasets of T cells sequenced from cancer patients treated with anti-PD-(L)1 ICB immunotherapies and examined evolution of T cell responses pre- and post-treatment. Quantitative of phenotypic and clonotypic entropy analysis with TCRi demonstrated improvements in characterization of the transcriptional signature of clonotypes. Furthermore, TCRi highlighted the importance of phenotypic flux and specific T-cell phenotypes as determinants of therapeutic response.

## Introduction

Human T cell repertoires have approximately 10^8^-10^10^ different clonotypes (Mora and Walczak 2019), each of which can potentially recognize a collection of antigens with varying specificity. Furthermore, individual T cells have a wide variety of phenotypes that determine their functionality. While this enormous diversity of clonotypes and phenotypes provides robust coverage against possible antigens, the response to viral infection or T cell dependent therapies is often driven by a relatively small subpopulation of the total repertoire. The identification and characterization of these subpopulations is essential to understanding the biological response but is experimentally and computationally challenging.

The ability to amplify and sequence the highly diverse T cell receptor (TCR) complementarity determining region (CDR3), which serves as a unique barcode for T cell clones and associated functionality, has led to quantification of statistics of the T cell repertoire by applying information theoretic diversity measures, such as Shannon entropy (Robins et al. 2009) to T cell receptor (TCR) sequences. Endogenous TCR sequencing has allowed the association of repertoire statistics with response to therapy (Leiserson et al. 2018; Tumeh et al. 2014), probabilistic models of TCR generation and selection (Sethna et al. 2019; Murugan et al. 2012), and motif-based TCR clustering methods that aim to address receptor specificity (Glanville et al. 2017; Dash et al. 2017). However, bulk TCR sequencing methods do not provide direct insight into T cell functional status and limit our ability to query whether antigen recognition, inferred by the narrowing of TCR clonotypes, is associated with specific phenotypes. Conversely, single cell transcriptomic analysis of T cells has led to the identification of transcriptional signatures and T cell subpopulations, which have been correlated with both response and resistance to immunotherapies (Caushi et al. 2021, Sade-Feldman et al. 2018). However, the consequences of these T cell subpopulations cannot be directly associated with antigen recognition in single cell RNA-seq alone. A more complete view of T cell functionality in the context of antigen recognition can be achieved only by combining both modalities.

Integrative measures bringing both TCR and phenotypic state into a unified description of the T cell have proved to be challenging (Valkiers et al. 2022; Zhang et al. 2021; Schattgen et al. 2021). Current methods have approached this problem in a variety of ways including the use of enrichment analysis within a set of phenotypes (Sturm et al. 2020), analysis of graphs that link clonotype and phenotype (Schattgen et al. 2021), and the generation of TCR-based embeddings (Zhang et al. 2021). These methods provide a clustering of clonotypes with an associated phenotypic state but rely on the discretization of T cell populations into clusters. Further, these tools focus on the constraint phenotype imposes on clusters of TCRs, with less emphasis on how antigen recognition may affect phenotype. In contrast, we introduce TCRinfo (TCRi), an informatic theoretic framework that combines probabilistic phenotype assignment with clonotype assignment in a joint probability distribution. TCRi naturally describes the two-way relationship of phenotype and clonotype, allowing the computation of three information theoretic metrics in a unified representation defined by three quantities: clonotypic entropy, phenotypic entropy, and phenotypic flux. Clonotypic entropy allows direct measurement of the diversity of clonotypes within a set of phenotypes. Inversely, phenotypic entropy provides a measure of entropy given the probabilistic assignment of the same set of phenotypes within a single clonotype. Finally, phenotypic flux measures the amount of phenotypic change within a clonotype over time, describing the changes induced by therapy.

We demonstrate the performance and insights from TCRi in four paired single cell RNA/TCR sequencing datasets derived from ICB treated cancer patients. Using our novel entropy-based measures, clonotypic entropy and phenotypic entropy (**Equation 1 & 2,** Methods), we quantitatively assess differences in response to anti-PD1 treatment across patients. We find lower clonotypic entropy in CD8 T cells with a higher probability of dysfunction, alongside both CD8 tissue resident memory (TRM) and CD8 precursor terminally exhausted (T_exp_) T cells that share dysfunctional gene expression features. Further, we observe that stability in the transcriptional signature of a clone before and after treatment, as measured by phenotypic flux (**Equation 3**, Methods), is indicative of a positive response to anti-PD1 therapy. Although intra-patient and cancer-specific signals are varied, our method provides a reproducible means of comparing results across existing and new datasets.

## Results

### TCRi framework

After determining the clonotype and phenotypic probability distribution of each individual cell, we construct the joint probability distribution of the T cell repertoire (**Figure 1**). We assigned a clonotype based on the nucleotide TRB CDR3 sequence and a phenotype, using methods form natural language processing (NLP) (Ceglia et al. 2022), from a set of predefined marker genes. However, other methods of defining clonotype (*c*) or phenotype (*f*) can easily be incorporated into our framework. The joint probability distribution of a repertoire, *p*(*c, f*), allows for easy restriction of analyses to specific subpopulations of the repertoire and information theoretic quantities to naturally characterize them.

**Figure 1:**
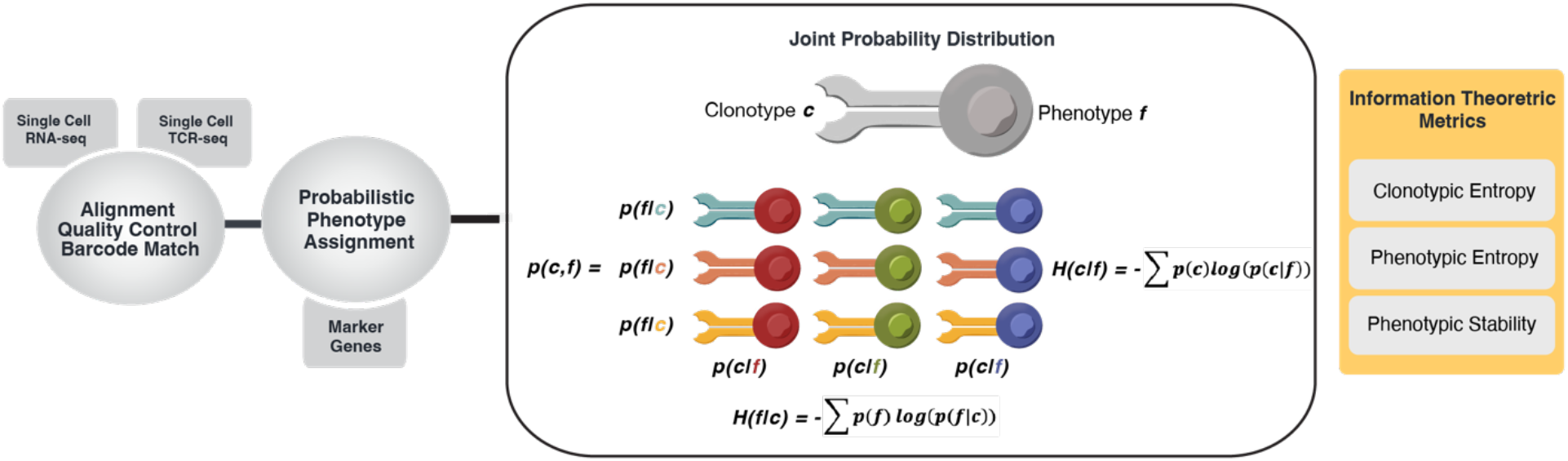
TCRi Framework and Joint Probability Distribution of Phenotype and Clonotype. Single cell TCR- and RNA-seq barcodes are aligned, filtered, and matched after sequencing. The gene expression is used to compute phenotypic probabilities from GeneVector using a set of marker genes. The phenotypic probabilities and TCR (TRB) sequences are taken as input into TCRi for the construction of the joint probability distribution *p*(*c,f*). Clonotypic entropy *H*(*c*|*f*) and phenotypic entropy *H*(*f*|*c*) re used to compute metrics and visualizations.

We define the notion of the phenotype distribution of a particular clonotype as the conditional distribution *p*(*f*|*c*), or similarly the clonotype distribution of a particular phenotype as *p*(*c*|*f*). These conditional distributions can be characterized by the diversity measures of clonotypic entropy of a phenotype and phenotypic entropy of a clonotype. Furthermore, if we have multiple samples from an individual, we can determine the changes between them by quantifying the differences between these distributions at each time point to yield a phenotypic (or clonotypic) flux using the L1 norm. In other words, the distance between two timepoints is assessed as the distance between probability distributions. Quantifying specific subpopulations provides better resolution than bulk TCR techniques, which operate on the clonotype marginal distribution *p*(*c*), or standard single cell or protein-based assays that operate on the phenotype marginal distribution *p*(*f*). We extended the framework to include a novel visualization of phenotypic flux as an alluvial diagram. The code is available as an open-source python library (https://github.com/nceglia/TCRi).

### Single cell analysis of anti-PD1 treated patients with carcinomas

We first examined 33,106 T cells with 28,371 unique TCR clonotypes in basal cell and squamous cell carcinomas (BCC, SCC) in 14 patients (Yost et al. 2019). We separated CD4 and CD8 T cells using GeneVector and subsequently classified four separate CD8 phenotypes and five CD4 phenotypes (**Figure 2A**) using a two-tiered analysis with a set of predefined gene markers (**Table 1,** Methods). CD8 T cells are classified into Dysfunctional, Activated, Naive, and Memory. Similarly, CD4 T cells are classified as activated, dysfunctional, T follicular helper (Tfh), T helper 17 (Th17), and naive. The top 10 clonotypes for CD8 T cells and CD4 T cells are densely clustered in the UMAP projection (**Figure 2B-C**) indicative of phenotypic similarity. We computed the phenotypic entropy, *p*(*f*|*c*), pre- and post-treatment for each patient (**Figure 2D**). We found that phenotypic entropy increases in 11 of 14 patients after therapy. Summarizing the phenotypic entropy across all patients, we find that this increase is statistically significant (p < 0.001) (**Figure 2E**).

**Figure 2:**
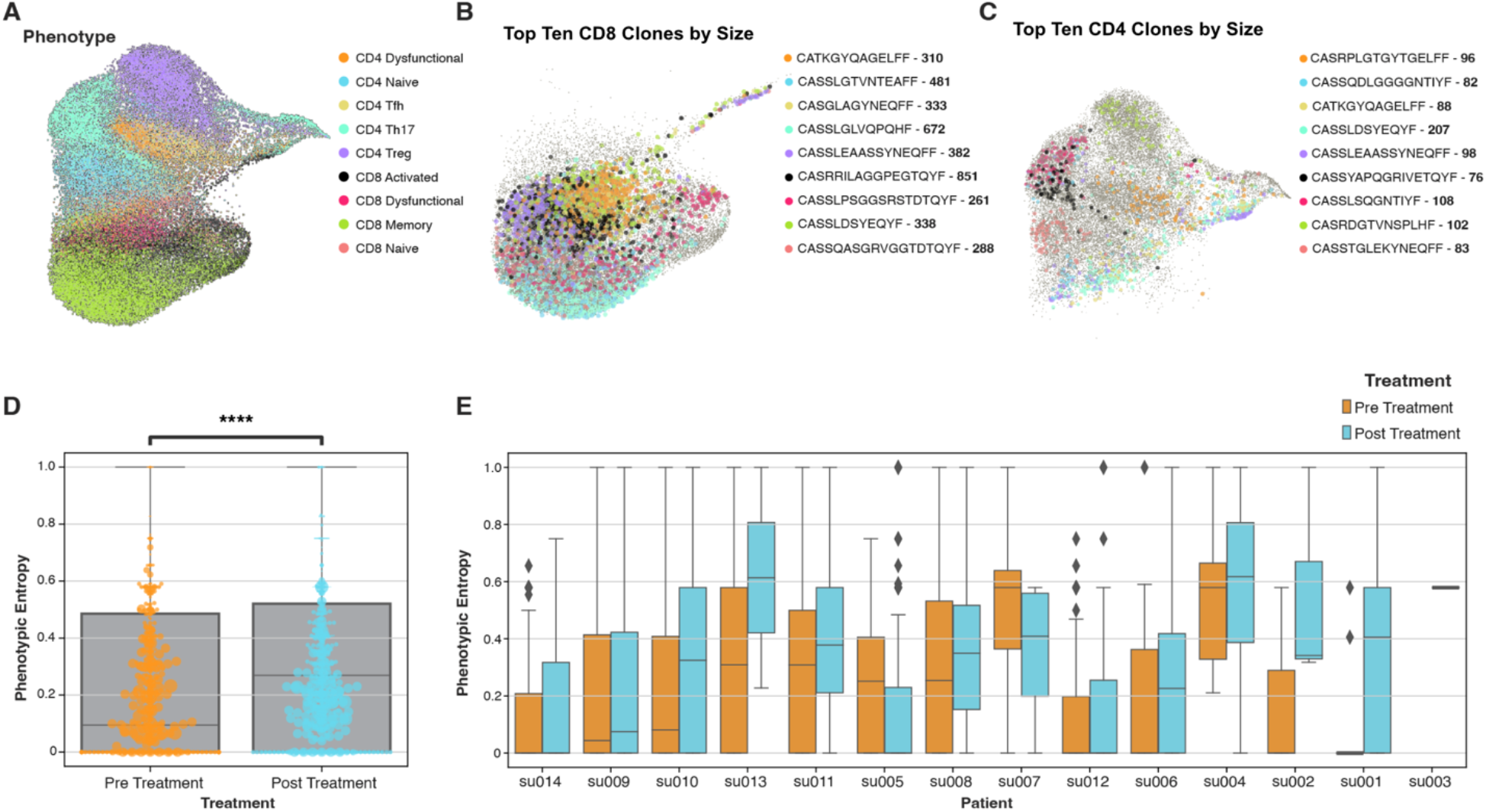
Probabilistic Assignment of Phenotype for T cells from patients with SCC and BCC. A) UMAP visualization CD4 and CD8 phenotypes for 12 patients for BCC/SCC dataset. B) The largest clonotypes within CD8 T cells placed in the UMAP. C) Largest clonotypes in CD4 T cells placed in the UMAP. D) Phenotypic entropy for each patient pre- and post-treatment (****, p < 0.001). E) Phenotypic entropy pre- and post-treatment summarized for all patients highlights a statistically significant increase in phenotypic entropy.

**Table 1:**
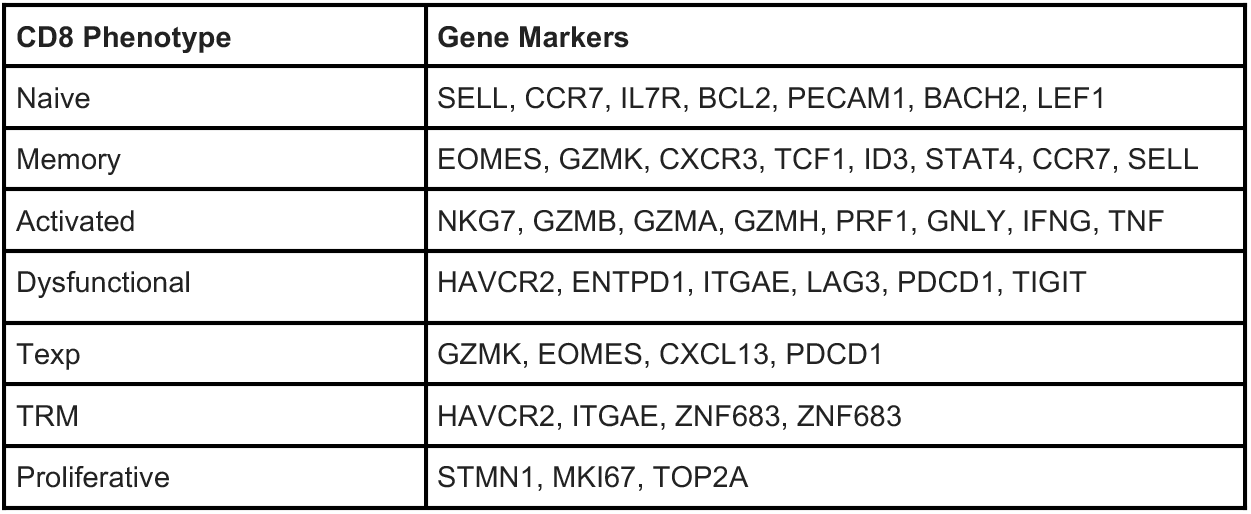
CD8 T Cell Phenotype Markers.

| CD8 Phenotype | Gene Markers |
| --- | --- |
| Naive | SELL, CCR7, IL7R, BCL2, PECAM1, BACH2, LEF1 |
| Memory | EOMES, GZMK, CXCR3, TCF1, ID3, STAT4, CCR7, SELL |
| Activated | NKG7, GZMB, GZMA, GZMH, PRF1, GNLY, IFNG, TNF |
| Dysfunctional | HAVCR2, ENTPD1, ITGAE, LAG3, PDCD1, TIGIT |
| Texp | GZMK, EOMES, CXCL13, PDCD1 |
| TRM | HAVCR2, ITGAE, ZNF683, ZNF683 |
| Proliferative | STMN1, MKI67, TOP2A |

Previous analysis of these data observed that as clones increase in size, the correlation of genes related to phenotype (cell type markers) also increases (Yost et al. 2019). This phenomenon was referred to as phenotypic stability. These large and phenotypically stable clones were found to correlate strongly with a dysfunctional T cell transcriptional signature. To investigate the relationship between T cell dysfunction and clonotype size, we computed the normalized clonotypic entropy for each phenotype from the joint probability distribution (**Figure 3A**). This measure is closely related to clonality using discrete phenotypes (1 - Normalized Entropy). By placing the measurement of clonality in the context of the probability of T cell dysfunction, we expand the utility of a familiar bulk metric to single cell analysis and provide a simple measure of clonal response in the context of treatment. Additionally, we observe that both naive CD8 and naive CD4 T cells have the highest normalized entropy, indicative of low levels of expansion; a finding that provides validation of our measure of clonotypic entropy.

**Figure 3:**
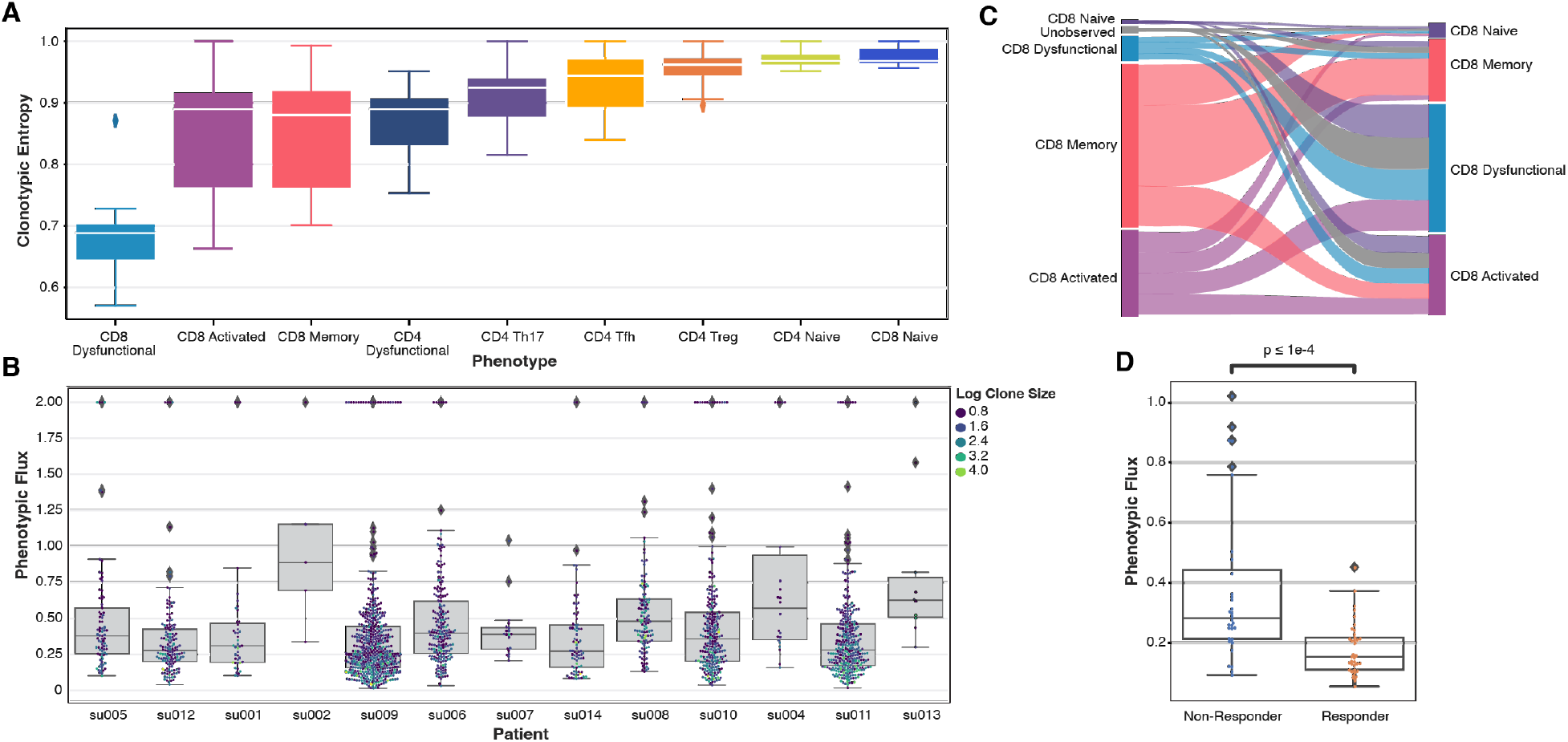
Information Theoretic Metrics of Clonal Changes in T cells from Patients with BCC and SCC. A) Clonotypic entropy for each phenotype over all patients. B) Phenotypic flux between pre- and post-treatment for each patient and colored by the number of cells in each clone. C) Probabilistic phenotypic changes normalized per patient for CD8 T cells including replacement clonotypes. D) Phenotypic flux quantified for non-responder and responder patients.

To investigate phenotypic flux as a measure of phenotypic stability, we computed the phenotypic flux (**Equation 6**, methods) of individual clones pre- and post-treatment within each patient. Here, we find that the largest clones have the lowest flux (**Figure 3B**). In other words, the transcriptional distance between pre- and post-treatment is smaller as clones increase in size. Describing the stability in terms of entropy provides advantages over identifying cell type markers correlate with large clones and a quantitative measure of stability as a small phenotypic flux. Specifically, we have shown that these large clones have an increase in transcriptionally stability beyond genes related to cell identity.

As phenotypes of individual T cells evolve in response to stimulation, it is critical to design metrics that explain these changes. Assigning hard classifications to individual clones by manual annotation of unsupervised clustering invariably produces expanded clonotypes with a spectrum of discrete phenotypes and no clear method to measure clonotypic changes. By incorporating the probability distribution of phenotypes, it is possible to generate measurements by which to quantitatively compare the phenotypic response within clones. Additionally, many new clonotypes can be identified after anti-PD-1 treatment that were unobserved before therapy. To generate a global map of probabilistic changes in phenotypes of CD8 and CD4 T cells, we summarized the likelihood of phenotype changes across patients, between pre- and post-treatment in the form of an alluvial diagram for CD8 phenotypes (**Figure 3C**) including unobserved clones. We find that the highest probable state for unobserved CD8 T cell clones is dysfunctional after treatment. To relate phenotypic flux to clinical outcome, we quantitatively assess the difference between clonotype repertoires at the pre- and post-treatment timepoints (**Figure 3D**). We identified a statistically significant difference in phenotypic flux between patients who responded to anti-PD-1 treatment, relative to those who did not respond. Specifically, we found that less flux is indicative of response to therapy.

These results highlight many of the original observations of this data and expand upon them to provide a more complete and quantitative description of transcriptional changes within T cell clones after anti-PD1 treatment. Specifically, we have shown that larger clones are more transcriptional stable, as measured by phenotypic entropy, and that this stability is evident across the transcriptome and not only restricted to dysfunctional markers. We have shwon stability is increased after anti-PD1 treatment (p < 0.001). Further, we observed that the CD8 T cell dysfunctional state as measured by clonotypic entropy, is the highlighted as least entropic, a finding that is closely related to clonality. Finally, we illustrate the transcriptional changes before and after therapy, highlighting those unobserved clones are most commonly dysfunctional and large phenotypic flux across T cell clones is indicative of clinical response. These results suggest that both phenotypic flux within the pre-existing clonotypes and the phenotype of additional infiltration of novel clonotypes after anti-PD-1 treatment, a phenomenon known as “replacement”, may contribute to the patient’s clinical response to therapy. While neither of these mechanisms are novel, quantifying this result directly in the joint probability distribution allows study to the effect of each mechanism in therapeutic response.

### Phenotypic Evolution and Clinical Response in Melanoma

Using TCRi, we processed 6,350 CD8 T cells in nine lesions from 34 melanoma patients treated with checkpoint inhibitors to examine changes related to clinical outcome in melanoma (Sade-Feldman et al. 2018). Using the same gene markers as input to GeneVector (**Table 1**, Methods), we find that all CD8 T cells fall into two main phenotypes: Memory and Dysfunctional. (**Figure 4A**). These results are nearly identical to those clusters previously reported (CD8_B and CD8_G), described as naive-memory and exhausted. Using the clinical response labels for each lesion, we measure the change in phenotypes for matched clonotypes using phenotypic flux (**Figure 4B**) and observe less phenotypic-clonotypic longitudinal change from pre- to post-treatment in patients responsive to therapy (p < 0.05), mirroring the trend in phenotypic flux found within carcinoma patients (**Figure 3D**). While these results only include CD8 T cells from two phenotypic states, the relative abundance of dysfunctional CD8 T cells is increased in patients with melanoma responding to immune checkpoint blockade.

**Figure 4:**
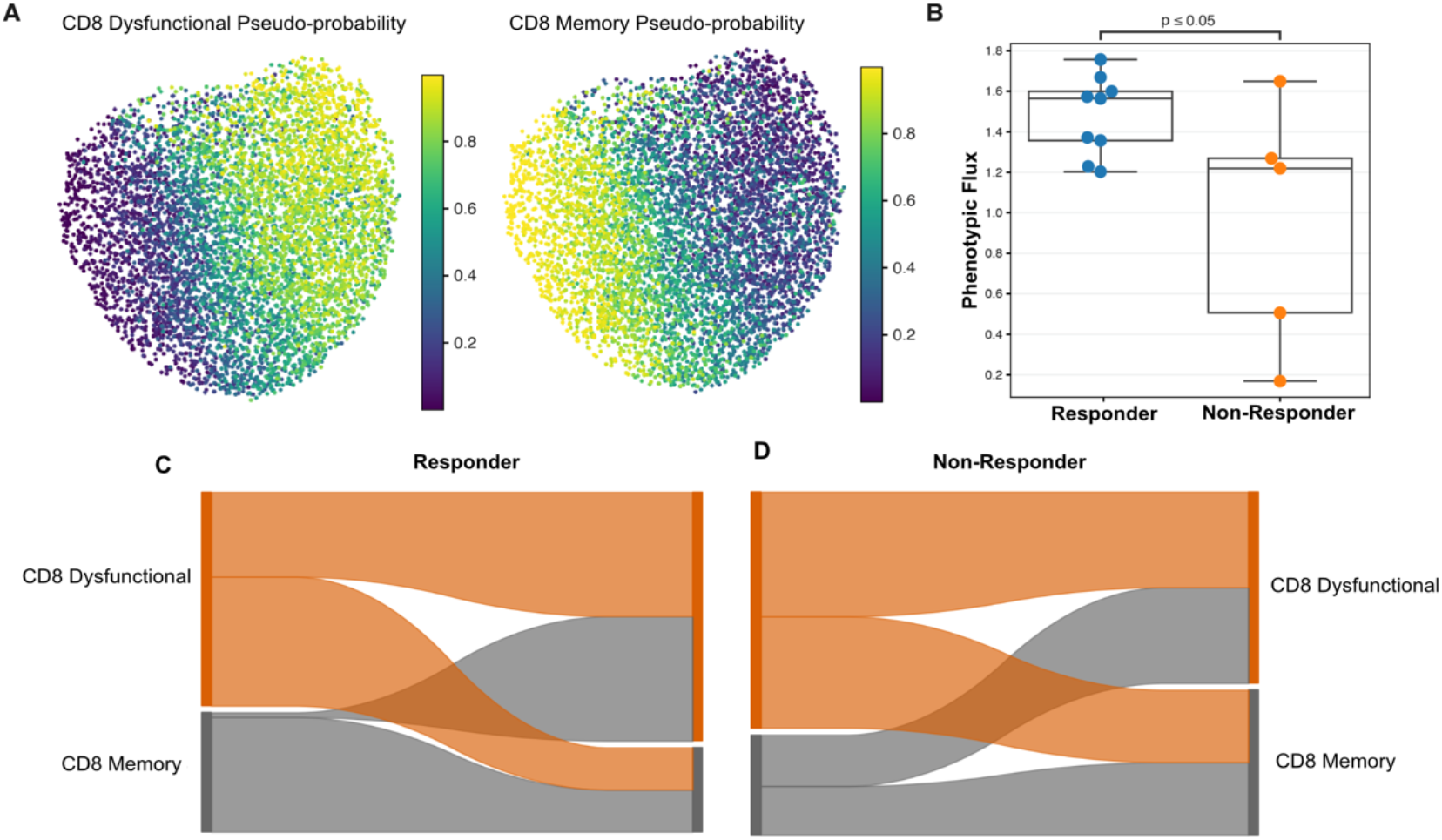
Clinical Response in Memory and Dysfunctional Melanoma CD8 T cells. A) UMAP of Dysfunctional and Memory phenotype probabilities that are representative of originally reported clusters but provide associated probabilities. B) Phenotypic flux of phenotype probabilities for patients with matched clonotypes pre- to post-treatment. C) Alluvial of phenotypic flux over treatment for responding patients highlighting that memory T cells retain phenotypic state after treatment and an increase in the proportion of dysfunctional T cells. D) Alluvial for responding patients shows higher phenotypic flux across both CD8 phenotypes.

After computing phenotypic flux, we visualized the probabilistic changes within clonotypes before and after therapy for non-responding and responding patients separately (**Figure 4C-D**) in alluvial diagrams. A dampened phenotypic flux in responding patients is most pronounced in memory T cells. Predominately, memory T cells in responding patients remain in a similar phenotypic state after anti-PD1 treatment. In contrast, many clonotypes from non-responding patients begin treatment with a larger probability of dysfunction T cells. These results again suggest that response to anti-PD-1 therapy can be identified in phenotypic flux between pre- and post-treatment samples, consistent in T cells across both squamous and basal cell carcinomas and melanomas.

### Identification of Partially Dysfunctional T cells in Anti-PD1 Treated Lung Cancer

The identification of phenotype states may be both improved and validated by accompanying clonotypes. In the case of precursor terminally exhausted T cells (T_exp_), previous methods have attempted to identify cells exhibiting these phenotypes by performing unsupervised clustering using TCRs that overlap with at least one initially identified dysfunctional cluster (Liu et al. 2021). This overlap can be efficiently summarized with phenotypic entropy using the probability of dysfunctional and T_exp_ states that share several exhaustion markers. Specifically, these T cells may exhibit both classic markers of dysfunction (TIGIT, ENTPD1, CXCL13, TOX, and HAVCR2) and simultaneously express pre-dysfunctional markers (GZMK, EOMES, and TCF1) (Van der Leun, Thommen, and Schumacher 2020). Equally as important, clonotypes that are found to phenotypically switch post-treatment to a terminally dysfunctional state with a higher probability are more likely to represent true T_exp_ cells.

Using 31,326 CD8 T cells from 8 patients with lung cancer pre- and post-treatment (Liu et al. 2021), we annotated six phenotypes, including a T_exp_ state (**Table 1**, Methods), and plot the probabilities for both the dysfunctional and T_exp_ phenotypes to demonstrate the distinct transcriptional signature for each phenotype in the UMAP embedding (**Figure 5A**). After classification of each of the six phenotypes (**Figure 5B**), we observed that the relative fraction of CD8 T_exp_ cells was increased at the post-treatment timepoint (**Figure 5C)**. Using clonotypic entropy between the dysfunctional and T_exp_ phenotypes (**Figure 5C**), we found that dysfunctional T cells have a significantly lower entropy than those T_exp_ cells (p < 0.01). Importantly, while the relative number of T_exp_ cells have increased, the lower clonotypic entropy highlights the transitionary state as compared with CD8 dysfunctional cells. Without combining both single cell gene expression and TCR sequencing, the increase in T_exp_ cells may be misinterpreted as a clonal expansion after therapy.

**Figure 5:**
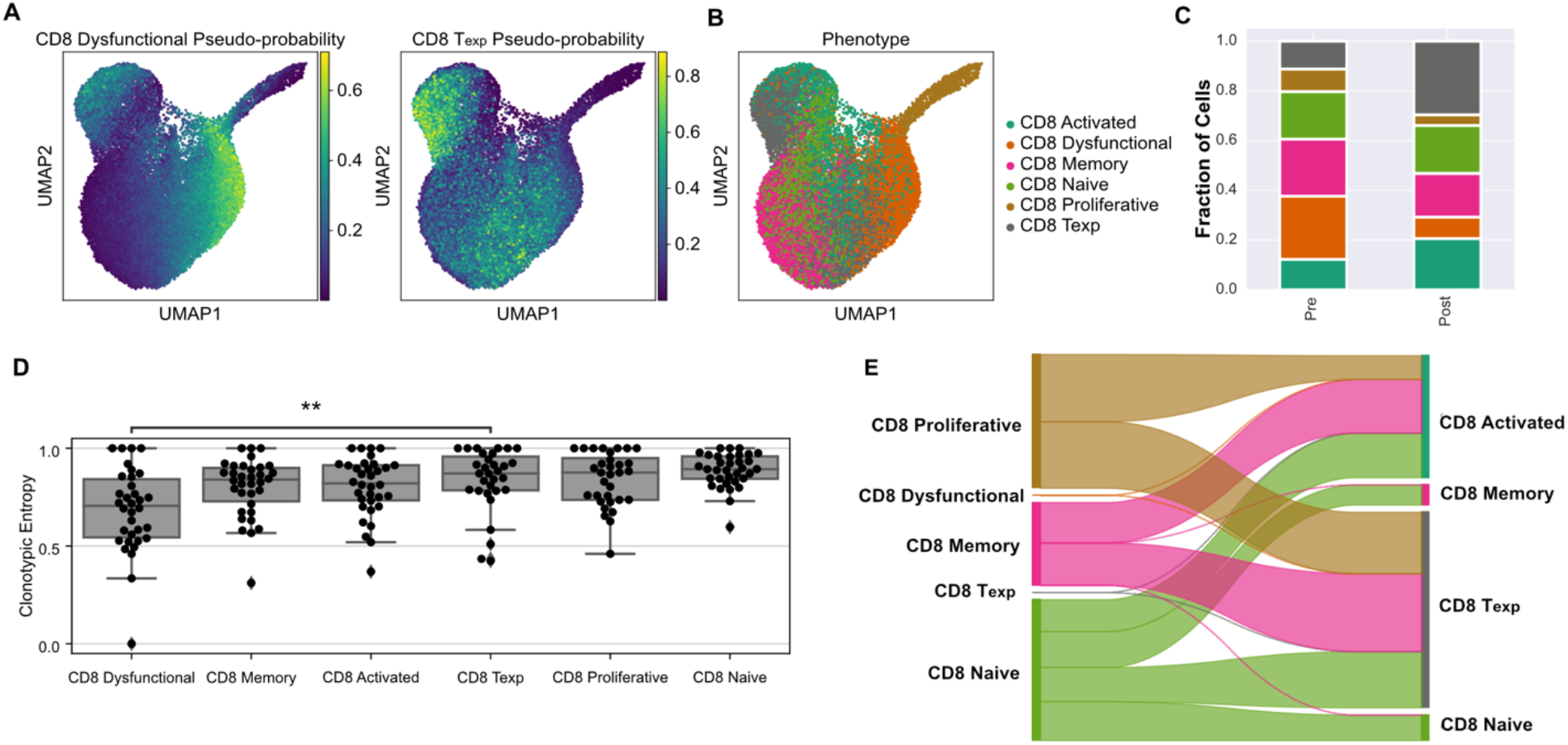
Identification of Precursor Terminally Exhausted CD8 T Cells in Lung Cancer. A) Dysfunctional and T_exp_ probabilities for each cell occupying transcriptionally separate space on UMAP. B) Classification of six phenotype states on the entire dataset. C) Compositional bar plot showing the expanded T_exp_ phenotype post-treatment (gray). D) Clonotypic entropy for CD8 T cells showing statistically significant difference CD8 dysfunctional and T_exp_. (**, p < 0.01) E) Alluvial diagram showing phenotypic flux for all phenotypes including dysfunctional and T_exp_ phenotypes.

In contrast, a lower clonotypic entropy reflects the narrowing the repertoire as T cells approach a terminal dysfunctional state, a result is also seen in carcinomas (**Figure 3A**). As T cells differentiate from a naïve state, the repertoire narrows, providing a clearer picture of the spectrum of CD8 T cell state lineage beyond cell state markers. We assessed that T_exp_ cells are both distinct from terminal dysfunction, where a larger number of clones have yet to expand, even if in higher proportion in post anti-PD1 treatment.

### Single Cell Analysis of Neoantigen-specific Clonotypes in Anti-PD1 Treated Lung Cancer

A quantitative description of the transcriptional landscape of T cells has the potential to identify mechanisms of resistance in the context of anti-PD-1 treatments. We have demonstrated the use of phenotypic and clonotypic entropy in measuring clonal expansion and the evolution of phenotypes for persistent and novel clones before and after PD-1 blockade. To highlight transcriptional signatures associated with known antigen specificity, we map phenotype probabilities from five transcriptional signatures (**Table 1,** Methods) to 235,851 CD8 T cells from nine patients with antigen reactive TCRs annotated from the MANAFEST assay (Caushi et al. 2021) (**Figure 6A**). Individual cells were labeled by major pathological response (MPR) for the patient of origin (4 MPR and 4 non-MPR) (**Figure 6B**). We computed the clonotypic entropy for each of the five phenotypes, and observe that alongside the dysfunctional and activated phenotypes, the tissue resident memory (TRM) signature exhibits low clonotypic entropy indicative of expanded clonal subpopulations (**Figure 6C**).

**Figure 6:**
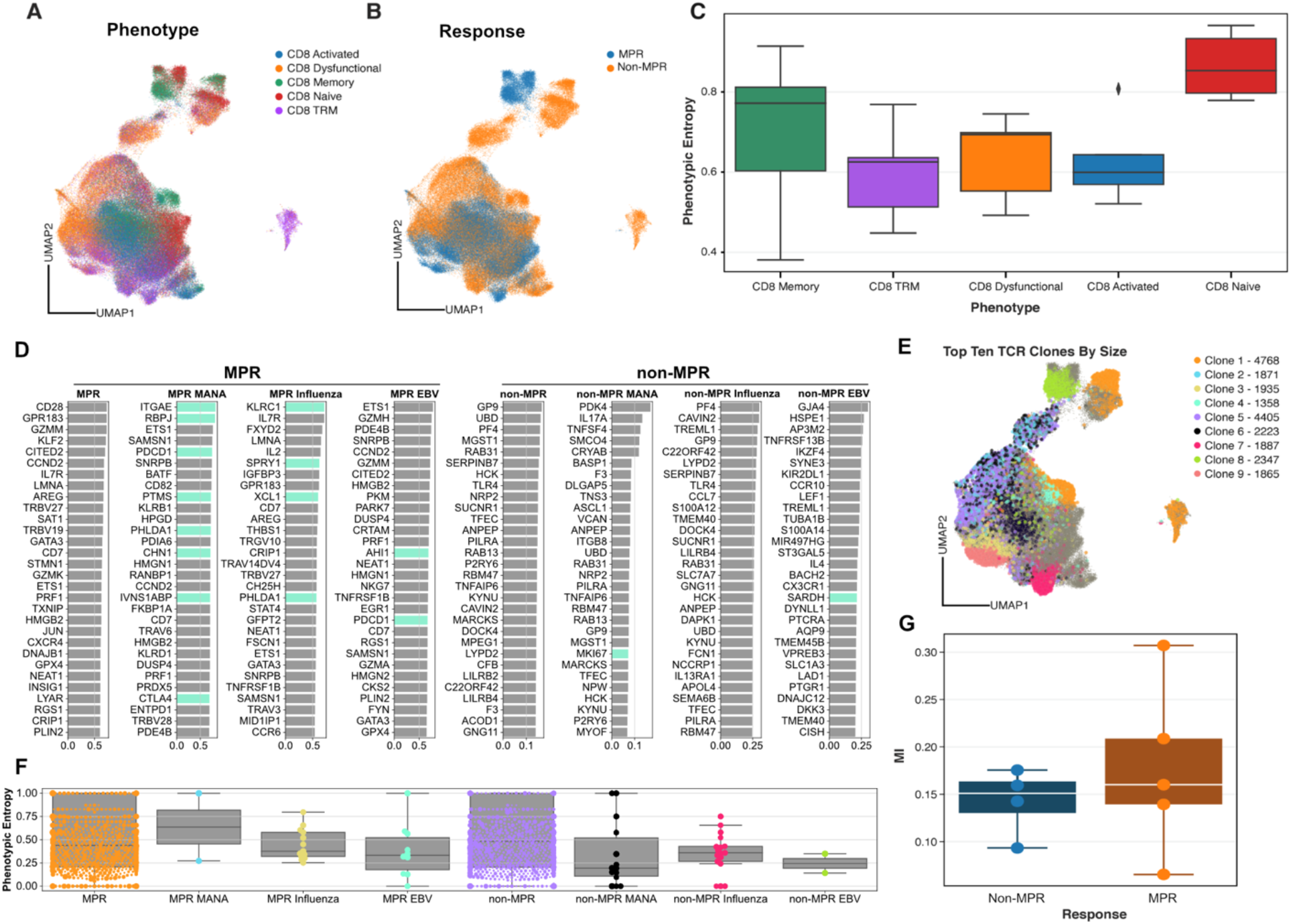
TRM markers and MANA-specific clonotypes. A) UMAP of phenotypes including TRM using gene markers as input to GeneVector (**Table 1**, Methods) B) UMAP plots showing major pathological response (MPR) versus non-responders (non-MPR). C) Clonotypic entropy over CD8 phenotypes highlighting TRM, Dysfunctional, and Activated CD8 phenotypic state with low entropic state. D) Most predictive genes for responders (MPR) and non-responders (non-MPR) for each MANAFEST annotated clonotype. T cells with MANA-specific clonotypes are most predictive of TRM associated genes (light blue). E) UMAP showing 10 largest clones including one with MANA specificity showing the variability in phenotypes for individual clonotypes when compared with A. F) Phenotypic entropy of antigen-reactive clonotype groups highlights the diversity of phenotypes existing within cells of identical clonotypes. G) Mutual information (MI) between clonotype and phenotype by patient per response.

To annotate antigen-specific clonotypes with unbiased transcriptional signatures, we compute linear vectors describing each clonotype using GeneVector (Ceglia et al. 2022). We identified the most similar genes to each of these vectors by cosine similarity in the lower dimensional space. We observe that responding MANA-specific clonotypes are most similar with TRM-associated genes (**Figure 6D**, light blue). These genes were identified via differential expression analysis (Caushi et al. 2021). We further examined the phenotypic entropy of each antigen-specific clonotype class separated by major pathological response (**Figure 6E**) and observed high phenotypic entropy in the MPR MANA clones with TRM signature. Our results indicate that antigen-reactive clonotypes are capable of existing and expanding in varying phenotypic states (as assessed by phenotypic entropy). Taking the top ten clonotypes by size (**Figure 6F**), we can visually assess this variability of phenotypes within individual clones on the UMAP embedding. While time series data was not available for this data, identifying the effect of clonotype on phenotype (as opposed to phenotype on clonotype) over time may prove important in understanding treatment response.

To further stratify MPR, we quantitatively assess patients by mutual information shared between clonotype and phenotype (**Figure 6G**). While there are not statistically significant differences between responding and non-responding patients, there is significant variability between individual patients, similar to the large patient variability seen in melanoma patients (**Figure 3C**). The computation of the joint probability distribution as a basis of TCRi provides a foundation for the additional information theoretic metrics to continue exploration of the relationship between phenotype and clonotype in cancer treatments.

## Discussion

TCRi provides a principled and quantitative framework, built on information theoretic measures, for describing clonotype and phenotype in single cell data. At the foundation, a joint probability distribution of clonotype and phenotype is constructed from probabilistic assignment of individual phenotypes. We demonstrate that two measures, clonotypic entropy and phenotypic entropy, provide an intuitive measure of the diversity of clonotypes within phenotypes, and vice versa. Additionally, we show that phenotypic flux, the phenotypic variability within a clonotype over time, can describe changes in phenotype distributions with treatment. Using these metrics, we visualized these change in novel alluvial diagrams.

Using four paired single cell TCR-seq and RNA-seq datasets, we highlight that these metrics quantitatively describe both novel and known mechanisms, including clonal replacement, of response to immunotherapy treatment. Specifically, we show with T cell data from patients with basal and squamous cell carcinoma, melanoma, and lung cancer that phenotypic entropy can demonstrate significant decreases in entropy after anti-PD1 treatment and, when paired with clonotypic entropy, can highlight high phenotypic variability in antigen-reactive clonotypes within phenotypes with clonal expansion. Additionally, clonotypic entropy can be used to further refine separate dysfunctional-like populations of interest, including T_exp_. Finally, that phenotypic flux can describe the stability of clones before and after treatment, and that this quantity may be indicative of response to anti-PD1 treatment. TCRi as a framework allows a principled view of the effect of phenotype on clonotype, and equally as important, on the effect of clonotype on phenotype.

While TCRi provides a foundation for quantitatively describing TCR-specific changes in single cell data, there are areas of improvement. Our metrics of entropy and flux have not uniformly separated clinical response variables across datasets. While this may be due to differences in types of clinical response assessments and small sample sizes, the addition of sub-phenotypes and pathways may prove useful in separating these patients. We envision our work can be expanded to atlas datasets to both annotate phenotypes and assess treatment in terms of clonotype reprogramming. Results may yet be improved by developing methodology to generate a joint probability distribution using graphical modeling, avoiding the need for empirical estimation, and improving computational speed.

While probabilistic phenotype assignment and entropy-based measures have been used previously in single cell RNA-seq and bulk TCR-seq studies, the novelty of our framework lies in their combination to describe paired single cell TCR datasets. Additionally, the reproducibility of these quantitative measures between patients is critical in the study of biologic response to immunotherapy at the single cell level, particularly as the use of immunotherapies continues to expand, novel therapies come online, and new combination therapies are put in use. By avoided ad-hoc clustering methods, TCRi provides a reproducible framework for testing gene signatures and deriving clonotypic entropy, phenotypic entropy, and phenotypic flux. TCRi creates a consistent mathematical framework for paired single cell data and, in doing so, can empower uniform comparisons of translational studies where T cell engagement is the key underlying mechanism of action.

## Methods

### Joint, Marginal, and Conditional Distributions

The joint phenotype-clonotype distribution can be constructed using either a probabilistic or deterministic assignment for either phenotype, *f*, or clonotype, *c*. Each cell can be assigned an individual joint distribution, *pi*(*f, c*), and the total joint distribution for a repertoire is easily computed by summing the distributions for each cell and renormalizing:

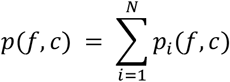

**Equation 1**: Joint probability distribution of phenotypes *F* and clonotypes *C.*

The initial phenotype assignment can be done in a variety of ways using whichever phenotype markers or method the user wishes if the total phenotype (or clonotype) assignment is appropriately normalized. For this paper, we curated a list of phenotype markers utilizing GeneVector in order to make deterministic and probabilistic assignments (Ceglia et al. 2022). The clonotype assignment is usually done deterministically by finding conserved positions on the DNA sequence of the TCR and extracting the CDR3 amino acid sequence. For this manuscript, a clone was defined by its CDR3 amino acid sequence, but our approach can also incorporate the V and J gene identities.

From the joint distribution, it is straightforward to compute marginal and conditional distributions. Previously, the marginal distributions, *p*(*c*) and *p*(*f*), have been studied and analyzed in a variety of contexts. The clone distribution of a single repertoire, *p*(*c*), is often used to compute the clonality/diversity of the repertoire or rank frequency distributions, whereas the phenotype distribution, *p*(*f*), is used to show the functional composition of a repertoire.

The conditional distributions are less often studied. The distribution of the phenotype of cells in each clonotype, *p*(*f*|*c*), describes the functional status of a specific TCR in the repertoire, while the distribution of clonotypes in each phenotype, *p*(*c*|*f*), can provide a more specific picture than the analyses of the clone distribution in the bulk (*p*(*c*)).

### Entropy

With our various distributions defined, it is useful to analyze the entropy of each distribution. First, the total entropy of the joint distribution is:

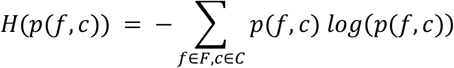

**Equation 2:** Total Entropy

The entropy of the marginal distributions, (i.e., the clone distribution and the phenotype distribution) are also readily defined by the *clonotypic entropy*:

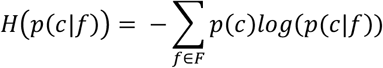

**Equation 3:** Clonotypic entropy for phenotype *f* over clonotype *c* ∈ *C*.

and the *phenotypic entropy*:

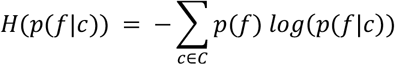

**Equation 4**: Phenotypic entropy of clonotype c over phenotypes *f* ∈ *F.*

The entropy of the clone distribution, *H*(*p*(*c*)), has been used many times before and is closely related to the normalized clonality measure used to characterize CDR3 distributions (Kirsch et al. 2015). The clonotypic and phenotypic entropies of the conditional distributions have not been studied before in this context. By defining the conditional distribution for these measures, we can also extend the notion of clonality to the conditional distributions and define the T cell receptor clonality of a specific phenotype. Lastly, as we have a joint probability distribution, *p*(*f, c*), we can compute the mutual information, *MI*(*f,c*), between clonotype and phenotype. This is given by:

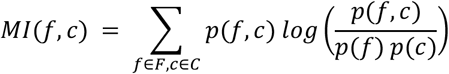

**Equation 5**: Mutual Information

### Comparing Repertoires

If we have two joint distributions, we can compare them (and the associated marginal and conditional distributions). We utilize the L1 norm:

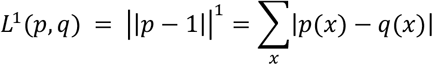

**Equation 6**: Phenotypic Flux: L1 Norm

This quantity is particularly useful when examining the conditional distribution of the phenotype of a given clone *p*(*f*|*c*). It can be used to show the changes of the clone between two different time points or locations. Large L1 norm can highlight a large amount of phenotypic reprogramming of the clonotype. Formally, we define the *phenotypic flux, L*^1^(*p,q*), as the L1 norm between the phenotype probability distribution in any two conditions. This quantity is most naturally used in measuring temporal change, such as before and after therapy.

### Phenotype Assignment

To formalize an annotation strategy that allows an unbiased assessment of perturbations, we propose a tiered classification strategy that first separates CD4 and CD8 T cells. These subsets are further classified into a broad set of phenotypes, in the case of CD8 T cells: Activated, Dysfunctional, Naive, and Memory. The markers used for phenotype assignment are based on previous literature and are used across all datasets in this study (van der Leun, Thommen, and Schumacher 2020; Akondy et al. 2017). We also note that within each subtype, we can selectively apply even more fine-grained phenotyping or pathway annotation including memory subsets such as tissue-resident memory T cells (TRM) and T_exp_ cells.

Within the GeneVector framework, a representative feature vector is computed for each marker gene set. Cosine similarity values for the given phenotypes are passed through a SoftMax function, given in Equation 6, to provide a pseudo-probability distribution for each phenotype. The argument maximum of this distribution is used to classify the most likely phenotype for a given cell.

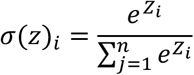

**Equation 8**: SoftMax function where *z* is the set of cosine similarities forn phenotypes.

### Predictive Marker Genes

Predictive genes for vaccine specific clones were generated by computing the low dimensional difference between vaccine specific and non-vaccine specific TCR+ T cells in the cell embedding and generating a sorted list of the most similar genes by distance in the gene embedding computed from GeneVector.

